# Cellbow: a robust customizable cell segmentation program

**DOI:** 10.1101/2020.04.21.052597

**Authors:** Huixia Ren, Mengdi Zhao, Bo Liu, Ruixiao Yao, Qi liu, Zhipeng Ren, Zirui Wu, Zongmao Gao, Xiaojing Yang, Chao Tang

## Abstract

Time-lapse live cell imaging of a growing cell population is routine in many biological investigations. A major challenge in imaging analysis is accurate segmentation, a process to define the boundaries of cells based on raw image data. Current segmentation methods relying on single boundary features have problems in robustness when dealing with inhomogeneous foci which invariably happens in cell population imaging. Here, we demonstrated that combined with multi-layer training set strategy, a neural-network-based algorithm Cellbow can achieve accurate and robust segmentation of cells in broad and general settings. It can also facilitate long-term tracking of cell growth and division. Furthermore, Cellbow is customizable and generalizable. It is broadly applicable to segmenting fluorescent images of diverse cell types with no further training needed. For bright-field images, only a small set of sample images of the specific cell type from the user may be needed for training. To facilitate the application of Cellbow, we provide a website on which one can online test the software, as well as an ImageJ plugin for the user to visualize the performance before software installation.

## 1. Introduction

Imaging has become a standard tool for the detection and analysis of cellular phenomena. Bright-field (BF) and fluorescent microscopy are widely used to quantify single-cell features^1^. The accurate quantification of such features critically depends on cell segmentation^2^.

Segmentation (the identification of cell boundaries for individual cells) is based on cell edge properties in images^3^. In fluorescent images, the edge properties of cells are very uniform, and only depend on the expression of fluorescent proteins (Fig. 1A). However, the typical appearance of a BF image depends on the imaging depth. As the depth changes, the images of cells change from bright border and dark interior to dark border and bright interior (Fig. 1B). Although this is often being used as an advantage to achieve the segmentation of cells, most of the existing methods rely solely on a single boundary feature^4–5^. However, due to the cell size variability and the imperfect alignment of cells with the focal plane, the problem of inhomogeneous focus often occurs^6^. Especially during cell growth, when the cell density changes rapidly, cells exhibit multiple edge features in the same image, e.g., when large cells exhibit bright edge features, small cells would exhibit dark edge features (Fig. 1C). As an algorithm based on a single feature would typically miss a subpopulation of cells, a large amount of subsequent manual correction work is required.

**Fig. 1.**
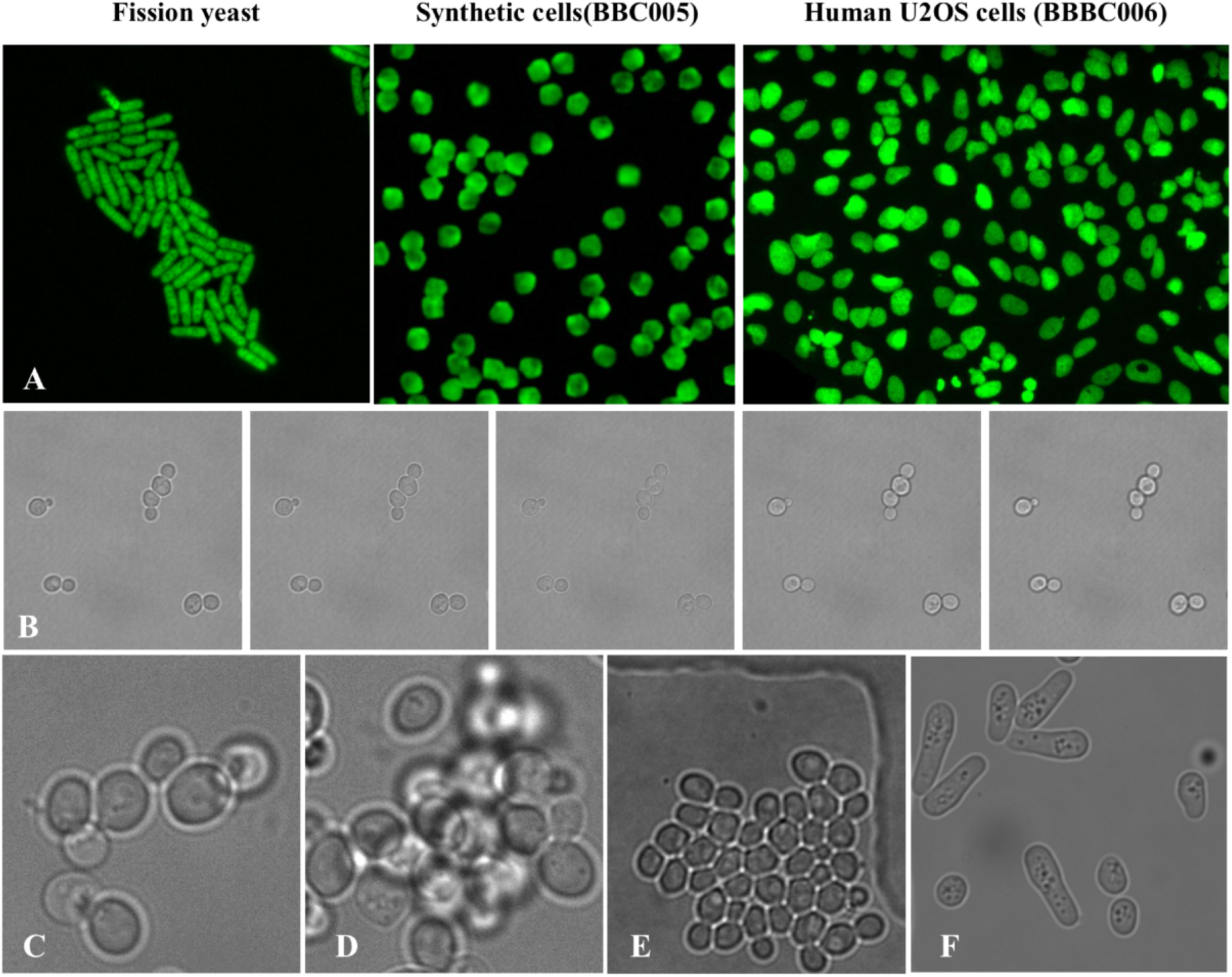
Most encountered problems in fluorescent and bright-field images. (A) Fluorescent images of differently shaped cells, the fission yeast images were from this study. The images of synthetic cells (BBC00S) and the human U20S cells (out of focus, BBBC006) were from the Broad Bioimage Benchmark Collection database. (B) The appearance of cells in a bright-field image depends on the imaging depth. As the depth changes, the images of cells change from a bright border and dark interior to a dark border and bright interior. (C) Inhomogeneous focus leads multiple edge features in the same image. **(D**, E) Agglomerated cells and culture dish edge exhibited similar local features to cells, even though they owned very different non-local features. **(F)** Rod-shaped and ball-shaped fission yeast cells.

In addition to local features such as dark or bright edges, cells also display many non-local features, such as specific shapes, size and length-to-width ratio. Such information is useful in identifying cells^3^. For example, floating agglomerated cells and impurities can exhibit edge characteristics similar to cells, but unlike cells, they have very different shapes (Fig. 1D, E). However, the discrimination of non-local features does not have a general solution^7^, so traditionally, different algorithms have been designed based on different cell shapes^8^. For example, algorithms for yeast cells are usually classified into either ball-shaped budding yeast algorithms^9,10,11^ or rod-shaped fission yeast algorithms^6,12,13^. In practice, we often need to integrate and discriminate many aspects of shape. For example, rod-shaped fission yeast appear spherical under certain culture conditions (Fig. 1F). Therefore, a universal algorithm for non-local feature recognition is needed.

Another common problem with the design of cell segmentation program was user friendliness. Although a large number of algorithms have been designed, they are rarely accessible to users. Users have to overcome the cumbersome steps of full software installation before determining whether or not the algorithm is useful for analyzing their own data. One solution would be that the algorithm designer provides users with an easy demo which is very convenient to test user’s own image, such as a website or familiar image processing platform like ImageJ^14^.

In the current study, we set out to develop a segmentation algorithm based on a deep neural network^15^ that can identify cell boundaries with inhomogeneous focus, using yeast cells as an example. It is a universal algorithm that can be applied to segment cells with multiple shape features and/or different imaging methods, such as ball-shaped budding yeast cells and rod-shaped fission yeast cells with bright-field images as well as fluorescent images. We then set out to design a website and ImageJ plugin for easy users’ test. Software for the algorithm is also available on the website.

## 2. Results

### 2.1 Multi-layer training dataset strategy solves the inhomogeneous foci problem

The difficulty of the inhomogeneous foci problem is that when the cells are at different imaging depths, their boundary characteristics will change (Fig. 1B). It could be solved by summarizing all the boundary features at various imaging depths, and carefully designing algorithms to identify them separately. This seemingly difficult task can be naturally accomplished by deep learning algorithm. Deep neural networks are good at extracting and summarizing boundaries features from the provided training images. Therefore, we trained the network to recognize multiple cell boundary features by providing a multi-layer training set.

We chose budding yeast to test the multi-layer training dataset strategy. Five layers of budding yeast images from 40 different fields of view, in which the cell boundary characteristics changed from bright border/dark interior to dark border/bright interior, were collected as the budding yeast dataset (Fig. 2A). Among that, 80% were used for training, 20% for testing. As the five layers of images were all from the same field of view, they shared a common labelling mask. Therefore, this strategy did not increase the annotation burden. As a control, we made a second layer of the 40 fields of view to provide a single-layer budding yeast training set in parallel.

**Fig. 2.**
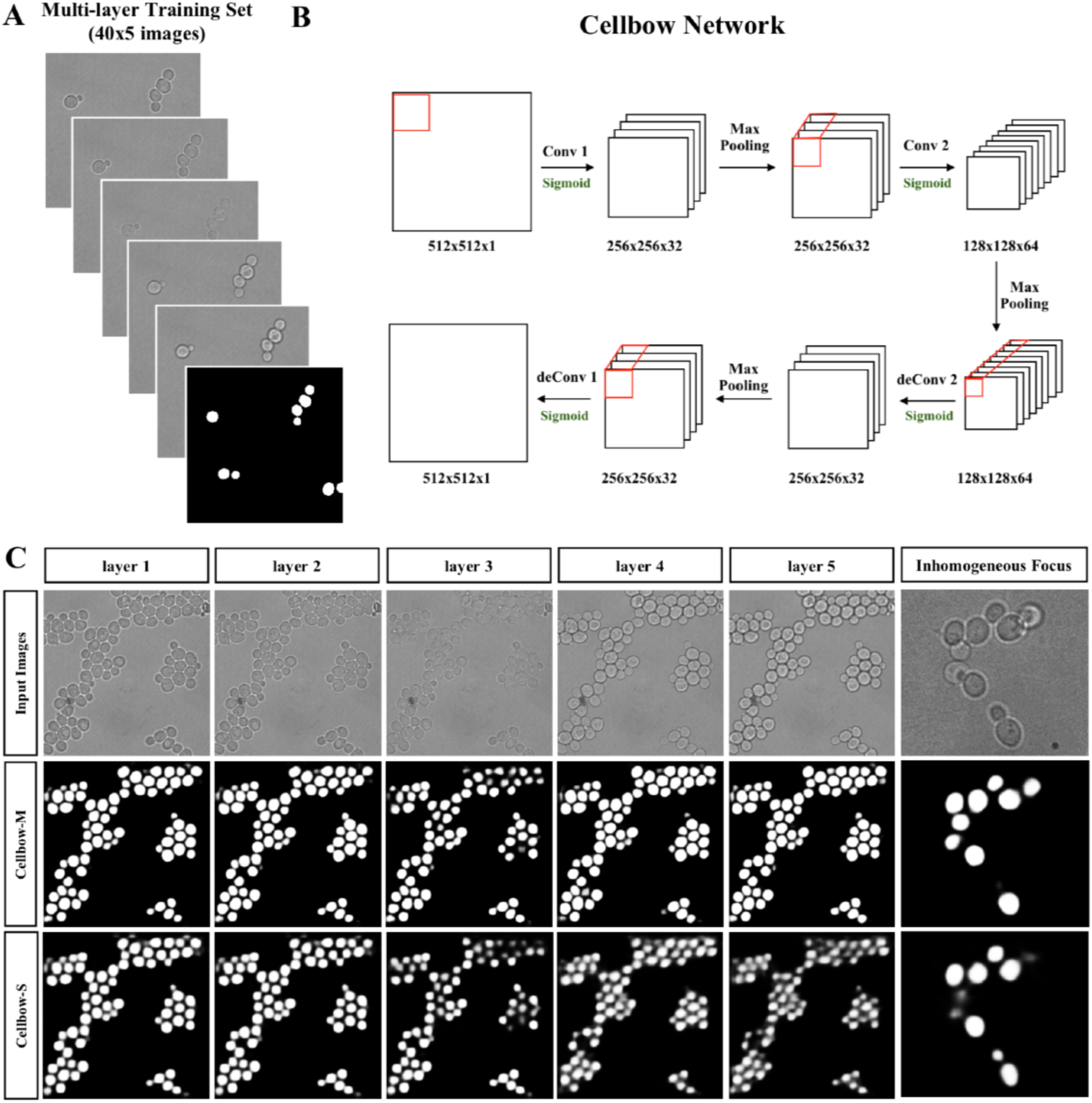
Multi-layer training dataset strategy solves the inhomogeneous focus problem. **(A)** Multi-layer training dataset included images from 5 different layers and one common labelling mask. **(B)** Cellbow architecture: for the coding part, there were two down-sampling convolutional and max pooling operators, and the de-coding part consists of two up-sampling de-convolutional and max pooling operators. Sigmoid was chosen as the activation function. **(C)** Top row: Test input image from 5 different layer and an inhomogenous focus test image. Middle row: Output mask from multi-layer training data et strategy (Cellbow-M). It successfully recognized cells from all five different layers. Simultaneously, it overcame the inhomogeneous focus problem. Bottom row: Output masks from single-layer training dataset strategy (Cellbow-S). It only recognized cells from the 2nd layer. However, it failed to overcome the inhomogeneous focu problem.

For the design of the neural network, we used fully connected neural network (FCNN) which has been applied to image segmentation tasks^16^ (Fig. 2B). The coding part, consisted of two down-sampling convolutional and max pooling operators, and the de-coding part consisted of two up-sampling de-convolutional and max pooling operators. The activation function was defined as sigmoid. Other detailed network structure parameters and training parameters are explained in the Methods. We name the network architecture “Cellbow”. After training, Cellbow was used to predict the cell body and background from a given new image. The predicted pixel values showed a bimodal distribution with peaks at 0 and 1, in which the background pixels were close to 0, and the cellular interior pixels were close to 1. Further thresholding was used to convert the prediction image into a binary mask.

Firstly, we demonstrated that the network trained with the multi-layer dataset strategy (Cellbow-M) successfully recognized cells from all five layers (Fig. 2C). Meanwhile the network based on the single-layer training dataset (Cellbow-S) only captured cells from the second layer (Fig. 2C). As expected, Cellbow-S failed to deal with the inhomogeneous focused cells. However, Cellbow-M captured both brighter and darker cell boundaries in the same image (Fig. 2C). Thus, the multi-layer training dataset strategy enabled Cellbow to overcome the most commonly encountered inhomogeneous focus problem during the imaging process, resulting in robust cell segmentation.

In order to quantify the prediction performance, we calculated the pixel-based F1, DI^17^ (Dice), and JI^17^ (Jaccard) based on the network prediction R and ground truth images S. The equations for F1, DI, and JI are given in the Methods. The average F1 of Cellbow-M was 0.93 (DI value, 0.93; JI value, 0.87).

### 2.2 Cellbow: Universal local and non-local feature extraction

To further investigate the ability of Cellbow to integrate and discriminate multiple cell shapes, we provided another multi-layer training set of bright-field rod-shaped fission yeast (Fig. 3A). In total, it contained 40 labeled focuses (each had 5 layers in depth). One hundred-eighty images were used for training and 20 were left for testing. Together with the ball-shaped budding yeast dataset, we retrained the neural network. Now named Cellbow-BF, it successfully identified both the rod-shaped fission yeast cells as well as ball-shaped budding yeast cells (Fig. 3B). In addition to the cell shape, we noted that it excluded floating agglomerated cells and culture medium edges which exhibited local characteristics similar to cells (Fig. 3C). This indicated that Cellbow was able to discriminate non-local features in the training set. The average F1 of Cellbow-BF was 0.87.

**Fig. 3.**
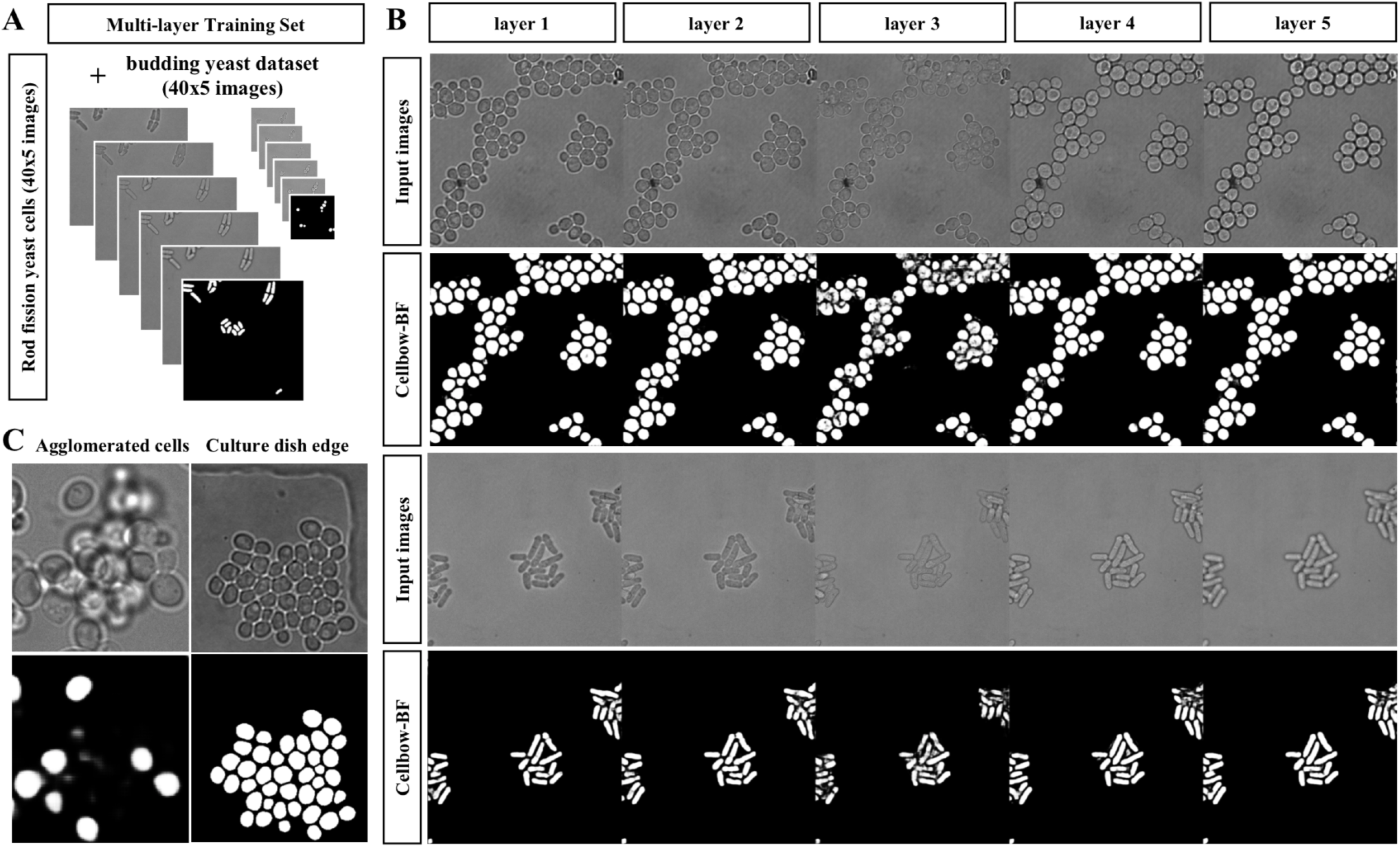
Cellbow can recognize multiple cell shapes in bright-field images. **(A)** Cell bow-BF was trained by using the multi-layer ball-shaped budding yeast and rod-shaped fission yeast dataset each included images from 5 different layers and one common labelling mask (each had 40 focus of view). **(B)** First row: Test budding input images from 5 different layer. Second row: Output masks from Cellbow-BF. Third row: Test:fission input images from 5 different layer. Forth row: Output mask from Cellbow-BF. **(C)** Agglomerated cells and culture dish edge input image (upper panels), and the prediction results from Cellbow-BF (lower panels).

### 2.3 Cellbow is universal and individually customizable

Cellbow was shown to be a rather universal algorithm that can summarize the local and non-local features in the training set. This would greatly improve cell recognition tasks. Traditionally, recognition algorithms were designed based on fixed boundary features of a given type images. Often different imaging methods and cell types used completely different algorithms. The user needed to search for a suitable software for his/her own project. This process was quite time consuming and energy exhausting.

Now the user can personalize Cellbow by offering their own training sets. Despite the fact that Cellbow can be a very accurate cell segmentation program, the required training set was small or even none. We demonstrated its versatility through the fluorescent image examples. In the fluorescent image, the cell boundaries only depend on the expression of fluorescent protein (Fig. 1A). Although the size and shape of different types of cells vary greatly, the feature of cell boundary is very consistent.

The training dataset contained 40 images of the fluorescence-labelled cytoplasm of fission yeast. We trained the same network as above, and named the trained network Cellbow-Fluo. We found that using only rod-shaped fission yeast as a training set, Cellbow-Fluo accurately segmented multiple cell types, such as the Synthetic cells (BBC005) and the Human U2OS cells (out of focus, BBBC0060) from Broad Bioimage Benchmark Collection^18^ (Fig. 4A).

**Fig 4.**
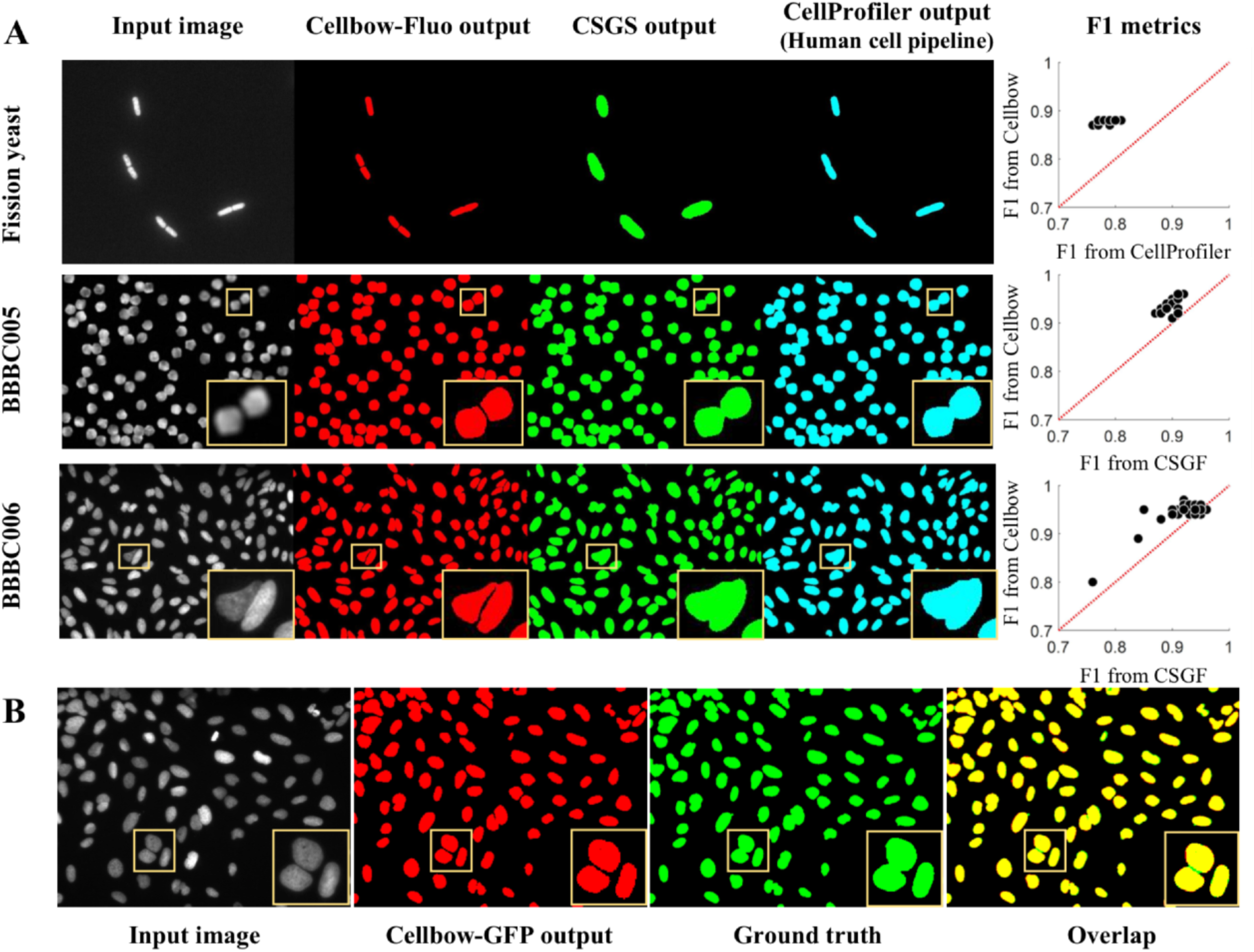
Fluorescent images of cells of various shapes can be segmentes by Cellbow. **(A)** Comparison of segmentation performance among different algorithms for different datasets. First column, input image; second column, output mask from Cellbow-Fluo; third column, output mask from CSGS algorithm; forth column, scatter plots of F1 metrics of the top two algorithms, each dot represented a test image. Top row: fission yeast cell fluorescent dataset; middle row: BBC005 datatset (synthetic cells); bottom row: BBBC006 datatset (out of focus, humans U20S cells). **(B)** Cellbow improved the segmentation of inter-cell gaps. From left to right: Input image, output from Cellbow-Fluo, ground truth image, and overlap of the output image and th ground truth image.

We compared Cellbow-Fluo with two fluorescent cell segmentation algorithms, the Cell Segmentation Generalized Framework (CSGF) ^19,20^ and the human cell pipeline in CellProfiler^5^. According to the ground truth provided by the database and segmented masks from three algorithms, F1, DI and JI were compared (Table1). The Cellbow-Fluo consistently outperformed the other two algorithms. This can also be seen from the scatter plots in Fig. 4A. We further analyzed where the accuracy has been improved. As it can be seen in Fig. 4A and B, the improvement of Cellbow was mainly located within the inter-cell gap, and these improvements were essential for accurate cell separation. In addition, we noticed that Cellbow’s differentiation of intercellular space was even better than the provided ground truth (Fig. 4B). Therefore, it can be seen that Cellbow not only achieved a significant improvement over the previous algorithms, but also needed no more training with further specific data for segmenting fluorescent images of diverse cell types. For bright field images, it may need training with a small set of customer-provided images.

**Table 1.** Comparison of segmentation performance among different datasets and algorithms.

|  | Fission yeast |  |  | Synthetic cells(BBC005) |  |  | Human U2OS cells (BBBC006) |  |  |
| --- | --- | --- | --- | --- | --- | --- | --- | --- | --- |
|  | Cellbow | CSGF | CellProfiler<br>(Human cell pipeline) | Cellbow | CSGF | CellProfiler<br>(Human cell pipeline) | Cellbow | CSGF | CellProfiler<br>(Human cell pipeline) |
| **F1** | 0.88 | 0.57 | 0.79 | 0.94 | 0.90 | 0.86 | 0.95 | 0.93 | 0.9 |
| **DI** | 0.88 | 0.57 | 0.79 | 0.94 | 0.89 | 0.86 | 0.93 | 0.91 | 0.89 |
| **JI** | 0.78 | 0.41 | 0.65 | 0.88 | 0.81 | 0.75 | 0.89 | 0.85 | 0.82 |

### 2.4 Accurate segmentation facilitates long-term monitoring of cell populations

Automated image analysis at the cellular level provides rich information. However, time-lapse cellular analysis is often hampered by inhomogeneous foci and the exponentially increasing cell density. In previous sections, we demonstrated that Cellbow, combined with a multi-layer training strategy, overcame the inhomogeneous foci problem robustly. To separate and identify single cells, we further applied distance transform-based watershed^21^ segmentation to the binary mask to achieve the final segmentation output (Fig. 5A). Once segmented into individual cells, we then identified the boundary, area, and centroid for each cell in the image. By using this algorithm, we tracked the cell number and cell size distribution of budding yeast and fission yeast (Fig. 5B, C).

**Fig. 5.**
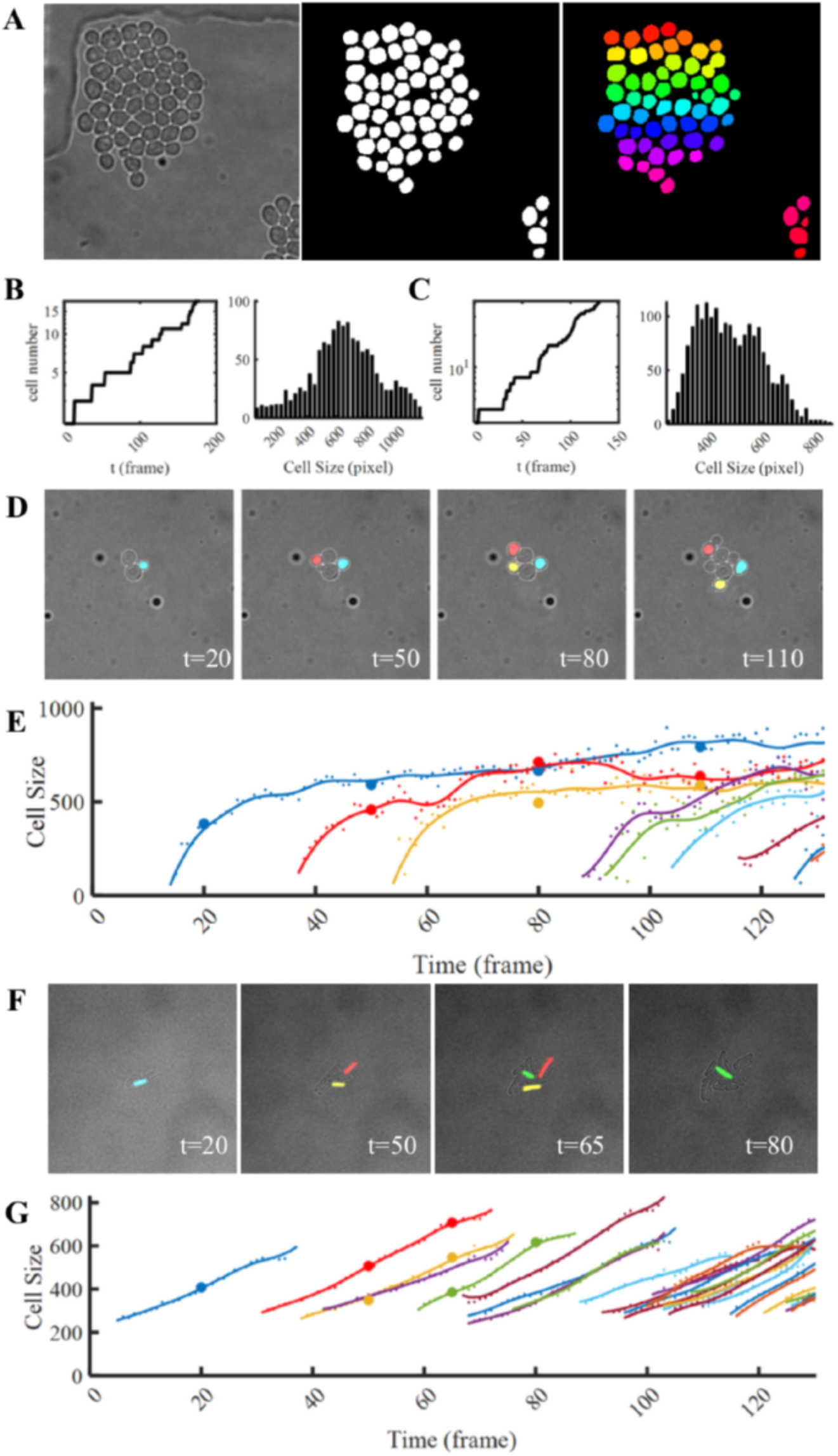
Cellbow enables long-term tracking of cells. **(A)** Input budding yeast bright-field image (left), output image from Cellbow-BF (middle) and final segmentation result (right). **(B)** Budding yeast cell number growth curve and cell size distribution. **(C)** Fission yeast cell number growth curve and cell size distribution. **(D)** Bright field images of budding yeast cells. Masks of three representative cells were shown. Four images represent time points 20, 50, 80 and 110. **(E)** Single budding yeast cell area growth curve. Three representative cells were marked with the corresponding colors. Time points 20, 50, 80 and 110 were labelled with thick dots. **(F)** Bright field images of fission yeast cells. Masks of four representative cells were shown. Four images represent time points 20, 50, 65 and 80. **(G)** Single budding yeast cell area growth curve. Four representative cell were marked with the corresponding colors. Time points 20, 50, 65 and 80 were labelled with thick dots.

To track the cells, we kept the cell body position in the image of the previous frame and searched for the most overlapped cell in the next frame. With this simple cell-tracking algorithm, we were able to trace the area growth curve of individual cells (Fig. 5D-G).

### 2.5 Cellbow Website

Users prefer to test their own images, but the cumbersome and time-consuming software installation steps deter many of them. To facilitate the adoption and future development of Cellbow, we set up a dedicated website (http://cls.pku.edu.cn:808/online/home/) and designed two demonstration versions and one full version of Cellbow. The demonstration versions were designed for the users to try their own data directly and quickly. It contained an online prediction website and an ImageJ plugin. The full version was tensor-flow-based source code^22^.

Website submission is easy and does not require any configuration by the user. A flowchart of how Cellbow predicts masks of cells from given images is shown in Fig. 6. The main webpages of the website are “Evaluation” and “Image Processing”. In “Evaluation” page, user can estimate the optimal objective magnification. It could be slightly different from the actual objective magnification value, because the performance of the network critically depend on the number of pixels occupied by a single cell. So the objective magnification difference depends on different imaging conditions and nutritional culture conditions. In “Image Processing” page, users can upload an image of their own, select the parameters, and click “Image Processing” button. Then, the cell mask images are generated and can be downloaded.

**Fig. 6.**
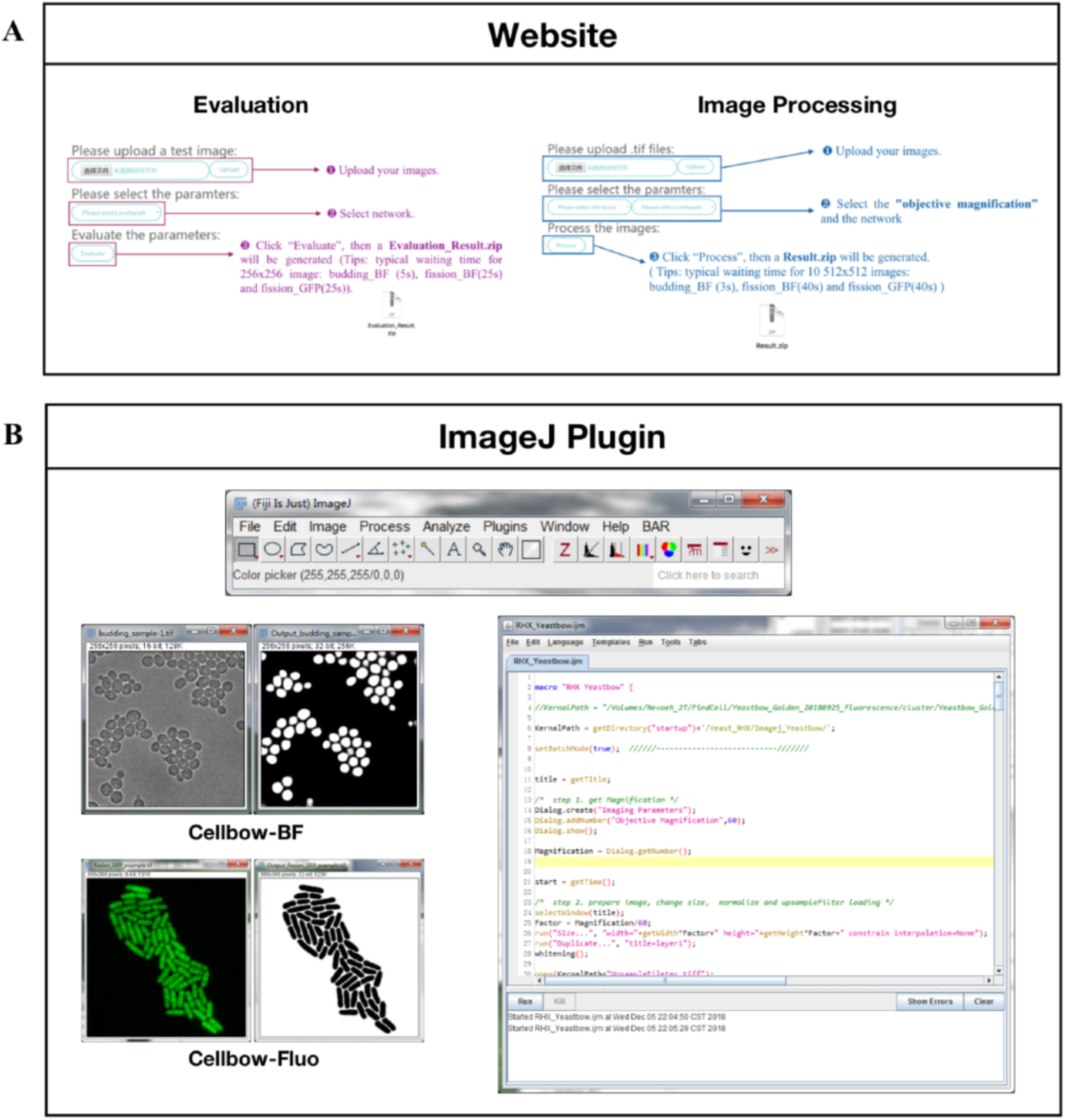
Website and ImageJ Plugin of Cellbow. **(A)** Two main flowchart on Cellbow website: “Evaluation” and “Image Processing”. **(B)** Two lmageJ plugins ar offered (Cellbow-BF and Cellbow-Fluo).

Another easy way to test Cellbow is using ImageJ plugin. Currently, we offer two plugins (Cellbow-BF for bright-field images of cells and Cellbow-Fluo for fluorescent images). Since it was written by macro language, no additional configuration is required. The user can just download the plugin and run it with their own image.

We strongly recommend the user use the website and/or the ImageJ plugin as a first step. After selecting the satisfactory version of Cellbow, they can apply the fully version. The software has very fast processing speed, e.g. it takes 1s for the segmentation of 10 images on a personal computer.

## 3. Discussion

In this work, we built a segmentation model Cellbow which simultaneously captured many features of cell boundaries in cell images. It overcame the most commonly-encountered inhomogeneous foci problem and facilitated long-term single-cell monitoring. Through the Cellbow website, users can test their input images following these steps:

1. For fluorescent images of diverse types of cells, user can upload their input images and get the output masks on the website (Cellbow-Fluo). Usually no custom training is needed.
2. For bright-field budding/fission yeast cells, users can upload their input images and get the output masks on the website (Cellbow-BF). Usually no custom training is needed.
3. For other types of images or when the user does not get a satisfactory results, one can personalize Cellbow with a labelling set of ~40 images.

Although in this article, we used multi-layer training set from the same field of view, but this is not necessary. Images can also come from different fields of view. As long as the training set contains multiple layers of data sets, the same improvement can be achieved. Compared with the two strategies, training sets from the same field of view reduces the labeling workload, and there is no essential difference otherwise.

When testing on the budding yeast, we noticed that the small buds of budding cells were sometimes missed by Cellbow. The main reason for this was that the manually labeled daughter cells in the training set were not perfect, and some smaller bud cells were omitted when they were manually labeled. Also, the daughter cells accounted for a small proportion in the training set, so they were biased. After discovering this problem, we perfected the labelling of daughter cells in the training set and retrained the network. Part of the daughter cells were identified, but there were still failed daughter cells. We need to work more in the future to solve this problem. This problem did not happen in the fission yeast cells. Therefore, the current algorithm was very successful for the statistics of mother cells, but caution should be taken when dealing with the budding daughters in budding yeast.

Finally, in applications we found that the follow-up segmentation and tracking procedure could be critical. Here we only used a simple watershed algorithm and centroid recognition to segment and track. In some cases, over-segmentation or under-segmentation can occur. Thus, for better performance it can be combined with some current downstream processing software.

## Acknowledgements

Part of the analysis was performed on the High Performance Computing Platform of the Center for Life Science.

## 4. Methods

### 4.1 Input Datasets

Training set generation is one of the most crucial steps for any neural network application. We first generated ground truth masks for the first layer. Then, the ground truth masks were generalized to the remaining layers which were acquired under the same focus of view. The filled cell body was chosen for the facilitation of final segmentation.

In this study, five input datasets were generated: budding yeast bright-field dataset (256×256 pixels), fission yeasts bright-field dataset (512×512 pixels), fission with various shape dataset (512×512 pixels), various contrast bright-field dataset (512×512 pixels), fluorescent dataset (512×512 pixels).

### 4.2 Image Preprocessing

This section mainly includes image labelling and augmentation step. The input images are regarded as matrixes of their original size square with labels in which inner-cell area is marked as 1 while background 0. In the process of data augmentation, we tend to acquire more images from origin sets to train the network with several ways like cropping, resizing and flipping, even though these new sub-images are exactly part of the original training set. However, they can actually provide efficient segment features to help the neural network achieve its best performance.

### 4.3 Deep Neural Network Architecture and Training

For encoding, we used two down-sampling convolutional and max-pooling operators; the down-sampling ratio was 1/2, take input image size of 256×256 pixels for example, it changed the image size from 256×256×1 to 128×128×16 in layer 2 and 64×64×32 in layer 3, and for decoding, we used two up-sampling de-convolutional and max-pooling operators. Notably, we chose sigmoid as the activation function following the convolutional and de-convolutional operators. The receptive field size of the FCNN was 5 in each layer, which is close to the diameter of a cell.

Several training hyperparameters are set as iteration steps = 100000 while learning rate = 0.0001 based on AdamOptimizer. After trained network predicting, one image for evaluating is transferred into a matrix **of** the same size while pixels are real numbers near 0 and 1, then the watershed algorithm is used to recognize independent cells. Our code is based on the open-source framework Tensor-flow, and trained on CLS H.P.C. (website: http://cls.pku.edu.cn:8080/clshpc/).

### 4.4 Segmentation and Post-processing

Yeast-bow network had a same input and output size, which realized pixel-to-pixel prediction. However, cells boundaries in the output probability mask may not be separated perfectly under the condition like cells from a high-density population or mother-daughter cells. To further separate and identify single cells, watershed segmentation was applied to the probability mask to get the final segmentation output. The input of watershed is a distance map, where the intensity of seeds has the lowest value. Finally, cell binary centers and minimal convex closure polygon boundaries are presented using another MatLab built-in function REGIONPROPS. Here, we ignore cells with an area less than a given threshold, here say it is 20 (default value).

First, only keeping the cells you want to track in the first image and erase the rest of the cells. Followed by iterative tracking. During each iteration, the position of the center of mass of the cells in the next image is first identified, and then it is determined whether each mass center is in the presence of cells in the previous image, and if so, the cells where the mass center is located are retained.

### 4.5 Evaluation Metrics

F1, DI (Dice Index) and JI (Jaccard Index) were used to evaluate the pixel-based segmentation performance of the FCNN by using an evaluation dataset. The prediction R from the network and ground truth images S determines these three metrics. The calculation of these metrics is given below.

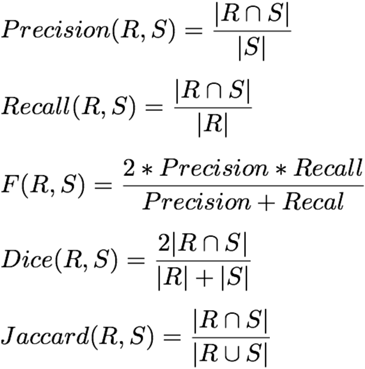

